# C-State: An interactive web app for simultaneous multi-gene visualization and comparative epigenetic pattern search

**DOI:** 10.1101/163634

**Authors:** Divya Tej Sowpati, Surabhi Srivastava, Jyotsna Dhawan, Rakesh K Mishra

## Abstract

**Background:** Comparative epigenomic analysis across multiple genes presents a bottleneck for bench biologists working with NGS data. Despite the development of standardized peak analysis algorithms, the identification of novel epigenetic patterns and their visualization across gene subsets remains a challenge.

**Results:** We developed a fast and interactive web app, C-State (Chromatin-State), to query and plot chromatin landscapes across multiple loci and cell types. C-State has an interactive, JavaScript-based graphical user interface and runs locally in modern web browsers that are pre-installed on all computers, thus eliminating the need for cumbersome data transfer, pre-processing and prior programming knowledge.

**Conclusions:** C-State is unique in its ability to extract and analyze multi-gene epigenetic information. It allows for powerful GUI-based pattern searching and visualization. We include a case study to demonstrate its potential for identifying user-defined epigenetic trends in context of gene expression profiles.

## Background

While the genome sequence of an organism remains fixed, different cell types exhibit characteristic and dynamic epigenomic profiles that lead to distinct transcriptional outcomes [1]. Information generated from next generation sequencing (NGS) following RNA isolation and ChIP (RNA-seq and ChIP-seq, respectively) is useful for understanding gene regulation. However experimental biologists often find it difficult to perform comparative analysis from large numbers of whole genome datasets. NGS pipelines encompass freely available and standardized algorithms to identify enrichment sites (reviewed in [2]), but bioinformatics proficiency is required to identify complex regulatory patterns. Apart from a few recent tools [3, 4], most pipelines do not support simultaneous analysis and visualization of data across genomic locations.

Querying gigabyte sized NGS datasets to highlight specific chromatin signatures thus remains a challenging task, requiring programming knowledge or familiarity with R (bioconductor) packages and working with the command line. Online tool portals such as Galaxy webserver [5] let users run command line tools by using a graphical front-end while the Integrative Genomics Viewer (IGV; [6]) needs to be installed on the users’ system. Most genome browsers typically allow linear sequence visualization but do not provide snapshots of epigenetic marks at multiple loci across cell types or experimental conditions, especially when these loci are distributed across the genome on multiple chromosomes. The UCSC genome browser [7] provides a powerful graphical user interface (GUI) to view user-specific as well as publicly available datasets and readily displays each genomic region at a high resolution. However, this platform presents difficulties in exploring multiple regions or genes when they are not linearly arranged along the chromosome, which is circumvented to an extent in the WashU browser [8].

Many of these browsers are webserver interaction based, maintaining server-side databases to generate web pages in response to user queries. This involves significant discontinuity in viewing large numbers of data points (genes and datasets), and time lost in data transfer. Traditional genome browsers relied upon client-server architecture due to limited client-side capabilities. However, the advent of HTML5 and several mature JavaScript frameworks in recent times allows the easy development of powerful interactive data analysis and visualization platforms that are independent of client-server interaction. The advantages to client-side rendering that overcome back and forth data transfer issues are outlined and implemented in JBrowse [9].

These features notwithstanding, most browsers are still not customized for the comparison of epigenetic patterns at gene subsets. Experimentalists currently have to deal with gigabyte-sized whole genome data and painstakingly extract the relevant information from tens of thousands of genes, followed by manually loading and examining each gene in succession. Viewing multiple selected loci in one go via a searchable GUI with continuous browsing ability thus remains a desired but largely unavailable feature.

Here we present C-State, a single-page application for comparative epigenomic analysis. Based on modern web technologies, C-State offers simple GUI based identification and comparison of epigenetic patterns at a large number of loci across multiple conditions and cell types. It provides a pipeline to load user-generated genome-wide peak information files as well as published ChIP-chip or ChIP-seq and RNA-seq datasets for simultaneous visualization and analysis of selected genes that may be located on different chromosomes. Its novelty lies in enabling interactive querying and filtering for enrichment patterns occurring within the selected target regions. Using C-State, epigenetic data trends can be easily compared with transcriptional profiles and plotted across all or filtered gene subsets to analyze the role of specific chromatin signatures.

## Implementation

### Resources

The genome information including gene and transcript coordinates, gene orientation, exon information, and gene description of various species and builds are downloaded from the Table browser of UCSC genome browser [7]. Wherever possible, multiple IDs of each gene are retained. The data are stored in tab separated flat text files, and a JSON file created to describe the one-to-one mapping of various gene IDs. Whenever a genome is selected, C-State refers to the JSON file to understand various columns of the tab separated file. C-State currently supports 14 genome builds of 8 species (see FAQs in website for full list).

### Program Architecture

C-State is a HTML5 based 100% client-side web app that can run on any modern browser such as Google Chrome (preferred), Mozilla Firefox, and Microsoft Edge. All algorithms of C-State, from data input to plot generation, are written in ES2015 (ECMAScript 6), the new standard of JavaScript. It follows the MVVM (model-view-view model) architecture based on VueJS (https://vuejs.org), and utilizes d3.js [10] to render plots in real-time. Object manipulation is handled by a combination of lodash (https://lodash.com) and custom functions. The modular architecture allows the customization of any aspect of C-State without affecting the functionality of other components. The collapsible accordions enable display of only relevant information, thus providing an uncluttered view of the task at hand while retaining easy access to the rest of the interface.

#### Script Usage and Logic

Users upload a simple text file containing the list of gene names/identifiers to be analyzed (Figure 1). Once the appropriate genome and build is selected by the user, C-State retrieves the corresponding genome information and uses the uploaded gene list to retrieve appropriate information such as the genomic locations, strand, exon information, and any neighboring /overlapping genes that map to the regions being analyzed. In cases of multiple transcripts, C-State considers the largest isoform. As genes can be located on either of the genomic strands, orientation of the gene must always be kept in mind during analysis. This causes visual discontinuity between gene upstream and downstream regions when comparing trends across multiple genes. To overcome this issue and enhance the visual similarity of genes on both genomic strands, all genes are corrected for their orientation and presented with a similar layout in the view panels.

**Figure 1:**
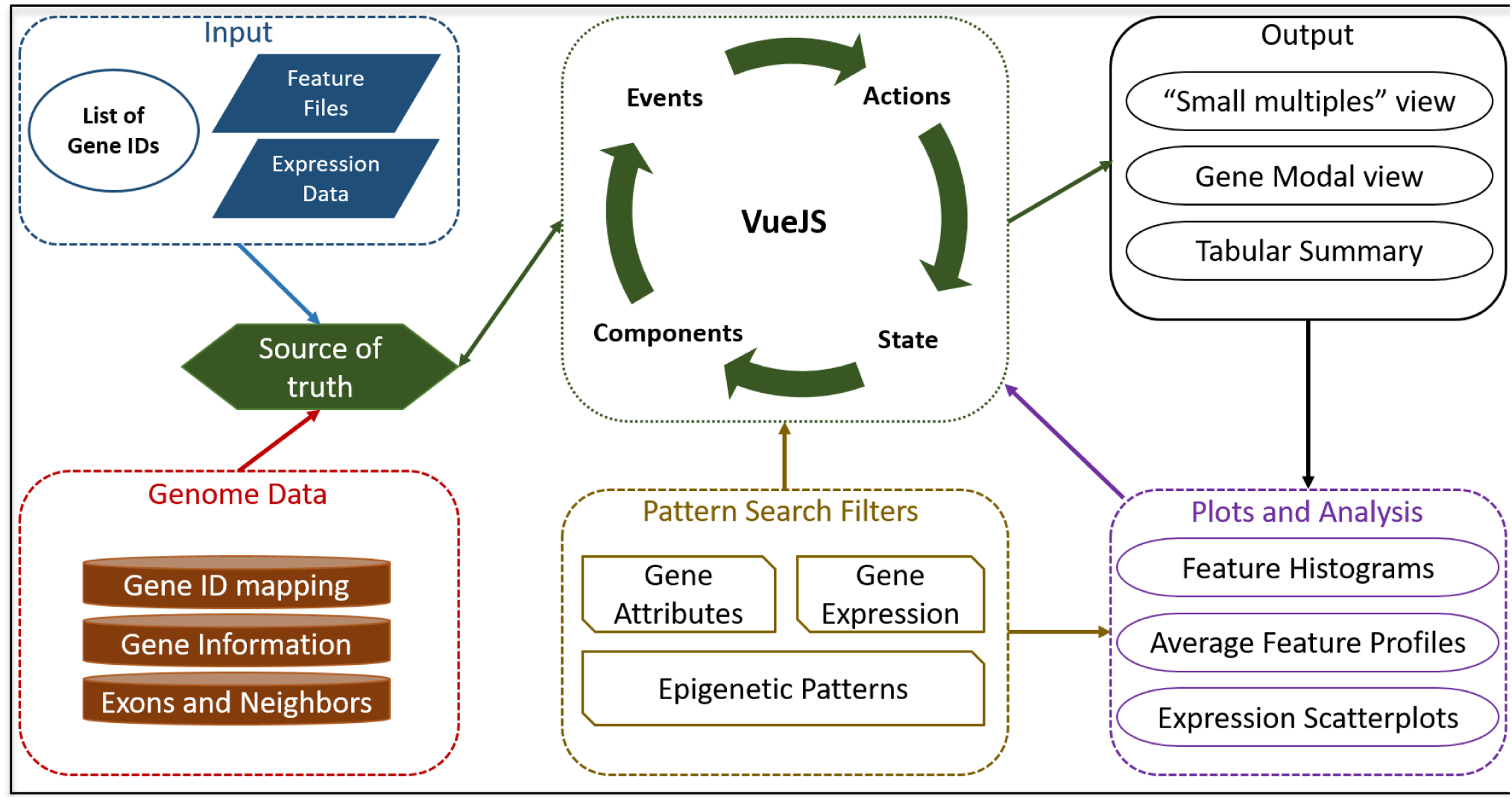
Overview of C-State’s architecture

**Table 1:** Comparison of C-State with popular genome browser

| C-State Feature | UCSC | IGV | WashU | JBrowse | CHROMATRA | CisGenome | Epiviz | visPIG |
| --- | --- | --- | --- | --- | --- | --- | --- | --- |
| Simultaneous display of multiple user-specified loci | ✗ | ✗ | ✗ | ✗ | ✗ | ✗ | ✗ | ✓ |
| Multiple features tracks | ✓ | ✓ | ✓ | ✓ | ✗ | ✓ | ✓ | ✓ |
| Concomitant display of gene expression data | ✓ | ✓ | ✓ | ✓ | ✗ | ✗ | ✓ | ✗ |
| Comparison of user-generated and published datasets | ✓ | ✓ | ✓ | ✓ | ✗ | ✓ | ✓ | ✓ |
| Versatile filters to allow selection of user defined patterns | ✗ | ✗ | ✗ | ✗ | ✗ | ✗ | ✓ | ✗ |
| Interactive plotting | ✓ | ✓ | ✓ | ✓ | ✗ | ✓ | ✓ | ✗<sup>5</sup> |
| Modular and Extensible | ✗ | ✗ | ✗ | ✓ | ✓ | ✗ | ✓ | ✗ |
| OS agnostic (works on multiple platforms) | ✓ | ✓ | ✓ | ✓ | NA<sup>3</sup> | ✓ | ✓ | ✓ |
| Programming knowledge not required | ✓ | ✓ | ✓ | ✗ | ✓ | ✓ | ✗ | ✓ |
| Prior dependencies not required | NA | ✓<sup>1</sup> | NA | ✗ | ✗ | ✗ | ✗ | ✗ |
| Runs locally | ✗ | ✓ | ✗ | ✓<sup>2</sup> | ✓<sup>4</sup> | ✓ | ✓ | ✓ |
| Open source license | ✓ | ✓ | ✓ | ✓ | ✓ | ✓ | ✓ | ✓ |

Key features of C-State checked for availability in other genome browsers and visualization tools.
✓: Feature available; ✗: feature unavailable; NA: not applicable
1 – Needs Java Run Time, which may not be preinstalled on some systems
2 – If using the JBrowse Desktop version
3 – Runs as a plugin for Galaxy Portal
4 – If Galaxy is installed as a local instance
5 – Plotting feature could not be tested on our data

To visualize gene features across the loaded genes, users can upload any number of feature files (histone peak information or other annotation files) and expression data files. C-State validates the file format to inform the user of any malformed lines, and calls its file reader function. Once all the files are parsed, the mapper function of C-State iteratively maps cell type/condition specific features and expression data to each gene of interest. Gene plots are rendered as SVGs using d3.js. The session information is stored in a single JavaScript Object, which can be downloaded as a JSON file that is self-contained and can be used in further sessions seamlessly. Each gene object exposes a Boolean property called “show” that can be used to toggle its display. This permits changing and customizing the genes to be displayed without modifying actual data.

### Design

As a 100% client-side app, C-State is designed from the grounds-up for high performance and efficiency. For example, analysis of data from 330 human genes and 24 feature files (4 histone marks from 6 cell types) has a peak usage of 1.1GB RAM, while loading 5000 genes uses 4GB RAM. VueJS is ideal for handling the view layer owing to its simplistic design choices, minimal overhead, and because it is non- opinionated about the underlying data structures. The UI is built with a minimalistic design using folding accordions for the Files (input) and View (output) panes, and a collapsible control panel for filtering and analysis. In order to resolve the issue of simultaneous multi-gene viewing across many cell types, the UI design utilizes the “small multiples” paradigm [11] where multiple genes are rendered as panels using a uniform co-ordinate system for visualization and comparison. C-State follows a component-based design; reusable logical structures are developed as individual components that can be incorporated anywhere in the app. This design is exemplified by the pattern search module of C-State where the filter layouts are separate components, and handle their logic independently of each other. This permits combining and chaining of numerous filters straight-forward, and each filter is contextually aware of the genes returned by all the previous filters. Further, as the filters can update the gene view simply by toggling the “show” property of any gene, the flow of logic is unidirectional and performance unaffected despite any number of genes or filters being active. The outcome of this granular design is a simple GUI-driven language using which the user can define a sequence of events (presence or absence of marks at specific locations) and C-State fetches instances where the definition holds true. This addresses the issue of complex pattern detection without resorting to coding.

Components of C-State are further organized as views, which are larger logical units. Communication between components is handled by a global event bus. To prevent misfiring, the event listeners are created only when a component is spawned, and destroyed as soon as the component is removed or obsolete. Gene panels are rendered as individual SVG charts once the event handler broadcasts to Vue that all the gene information is ready. Whenever a gene header is clicked, the modal view handler is populated with the appropriate data, and triggers the updated view. Linked zoom of all data tracks in modal view is achieved by broadcasting all mouse events with the x, y, and *scale* values as payload. Other data tracks listen to these events and update their own values accordingly.

Since plots in C-State are SVG containers, updating the view may require redrawing several thousand SVG nodes. As update events are asynchronous, requesting all elements to be redrawn simultaneously can throttle the CPU resources and may crash the web browser. C-State handles this by introducing a small imperceptible delay in firing the redraw events. The delay is calculated dynamically based on gene size and the indices of cell type and features, and is enough to permit the CPU to finish any pending operations.

## Results

### Overview

C-State provides an epigenetic pattern search and query platform for gene-centric analysis across a large number of loci. It retrieves co-ordinate information for user-defined genes and the genomic regions around them from whole genome datasets that are normally tedious for non-bioinformaticians to handle. The interactive and user-friendly GUI filters and displays the loci of interest using multiple criteria without the need for any computational knowledge. The input for the application is a simple list of gene names or identifiers (IDs) and ChIP-seq and RNA-seq datasets. By eliminating the need for any pre-processing, data transfer or installation, C-State gives biologists direct access to genome-wide data on their desktop devices for epigenetic analysis and biological interpretation.

The following sections describe the workflow for the analysis of epigenetic features in the context of varying gene expression status in different cell types.

### Data import: Files accordion

C-State primarily requires a list of genes for which the epigenetic data is to be analyzed along with the genome and build information (Figure 2). Users can specify whether they are interested in information only from the gene body or require flanking genomic regions around the target gene. The selection applies across all analysis and visualization modules and is set to 20 kb upstream and downstream of each gene in the genes list by default. C-State provides flexibility on the go in selecting target regions for analysis; the flanking regions can be changed using the flank selector and C-State reanalyzes the plots. This feature is useful in identifying epigenetic patterns underlying putative gene regulatory elements, which often lie in gene-proximal intergenic regions.

**Figure 2:**
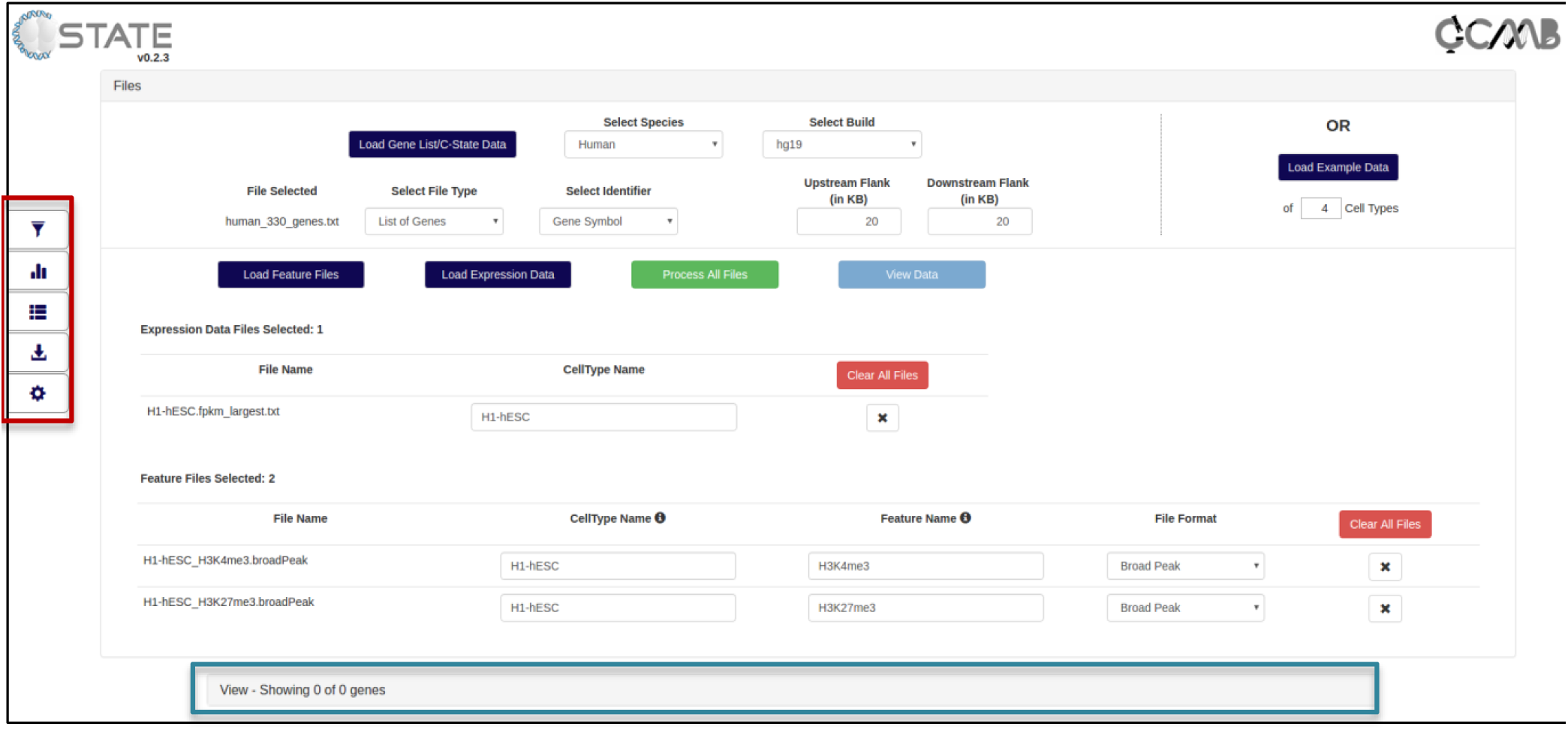
Open Files accordion of C-State showing data import and auto-mapping of the file attributes. The collapsed View accordion is indicated with a blue rectangle and the control panel is highlighted with a red rectangle.

C-State directly accepts genome-wide enrichment data (Features) files and, optionally, the expression data for each of the chosen cell types or experimental conditions. The features files constitute the peak information as obtained from a public database or generated from the user’s experiment (any genome coordinate-based information can be input as feature files including, but not limited to, ChIP-seq datasets, CpG islands, DNase hypersensitive sites, restriction enzyme sites and repeats). C-State accepts the widely used BED, broadPeak and narrowPeak formats for input of genome-wide datasets. File attributes are auto-mapped for plot generation based on the file names (Figure 2) and can be modified if needed.

### Control panel

Downstream functionalities of C-State such as the ability to identify epigenetic patterns and analyze genes bearing selected features are accessed from the 5 control panel keys on the left (Red box in Figure 2).

#### Pattern search module

A distinct feature of C-State as compared to traditional genome-browsers is its search and filter functionality for analyzing patterns in the input data. Simple filtering is based on identifying regions specified with a set of operators appearing from a drop down menu in filters for gene name, size, chromosomal and genomic context, transcript levels and presence or absence of peaks at user-specified locations (Supplementary Figure S1).

C-State also provides the ability to build complex queries using simple text by chaining multiple filters in any order. This helps the user define conditions to look for specific relationships between any pair of features, for instance overlapping peaks indicating bivalent domains (see use cases below) or peaks juxtaposed upstream or downstream to each other within a specified distance. Searches can be refined by specifying the distance of the pattern from the TSS as well as the cell type. The number of genes filtered out at each stage is displayed on the Filter and the total number of genes is indicated in the View pane, above the legend.

#### Plots and analysis

Clicking this button takes the user to the plots area to analyze global trends across cell types. These include feature histograms and feature profiles with respect to TSS or gene bodies as well as gene expression scatterplots. Activating the pattern search filters in the previous module enables data plotting only from filtered gene subsets carrying a specified epigenetic pattern for comparison with the whole genome trends.

#### Tables

C-State provides an interactive format for tabulating information of all loci in the input list or of filtered genes in the gene set that share a common epigenetic trend. The table is linked with the gene modal view for visualizing locus-specific data as described in the next section. Gene and gene expression details are also available in the table. The gene names or IDs can be directly copied from the table and saved for further GO or other analysis.

#### Downloads

This tab provides three download options: i) a text file that summarizes the analysis performed, including any filters that have been set and the list of genes that pass the filters, ii) a single SVG file of all the genes that are currently displayed - this file can be further edited in other image processing software to generate high quality images and iii) a fully-functional self-contained JSON file that can be uploaded to C-State directly to re-initiate the session and continue the analysis. The JSON file can also be shared with collaborators, allowing them to view and analyze data without the need for sharing any raw data files.

#### Settings

The settings menu provides the user with options to customize various aspects of C-State; for instance, peak score and size cutoffs can be changed in order to analyze only high quality peaks in the datasets. C-State also provides for extensive customization of the view panels (toggle display of neighboring genes and exons), feature tracks and expression data scale, and color schemes.

### Data output and visualization: View accordion

To convert the files into input files for the display module, C-State parses the genome-wide chromatin and expression datasets to retain only the features relevant to the user’s interest based on the genes/regions list provided (elaborated in the Implementation section). Following a gene-centric approach, the genomic coordinates specified using the BED format are converted relative to the transcription start site (TSS) of each gene. All the peak features and genes are then corrected and re-plotted with respect to the TSS, thereby allowing a more intuitive and direct comparison of genes on both positive and negative strands of the genome.

#### Default display

On loading the data files, the View accordion opens and the Visualization pane gets populated with gene-specific data panels, arranged based on the number of conditions / cell types loaded. The number of genes displayed is indicated along with a legend for the feature and expression tracks loaded (Figure 3A). A quick search bar allows for rapid browsing of specific gene(s). The data of multiple cell types is arranged column-wise (labelled at the top of the column); data for each gene is thus displayed side by side across all samples under consideration, facilitating comparative visualization. The visualization pane uses dynamic width for plots and adapts to the number of cell types/conditions uploaded, so that the plots are not rendered off-screen (Figure 3B).

**Figure 3:**
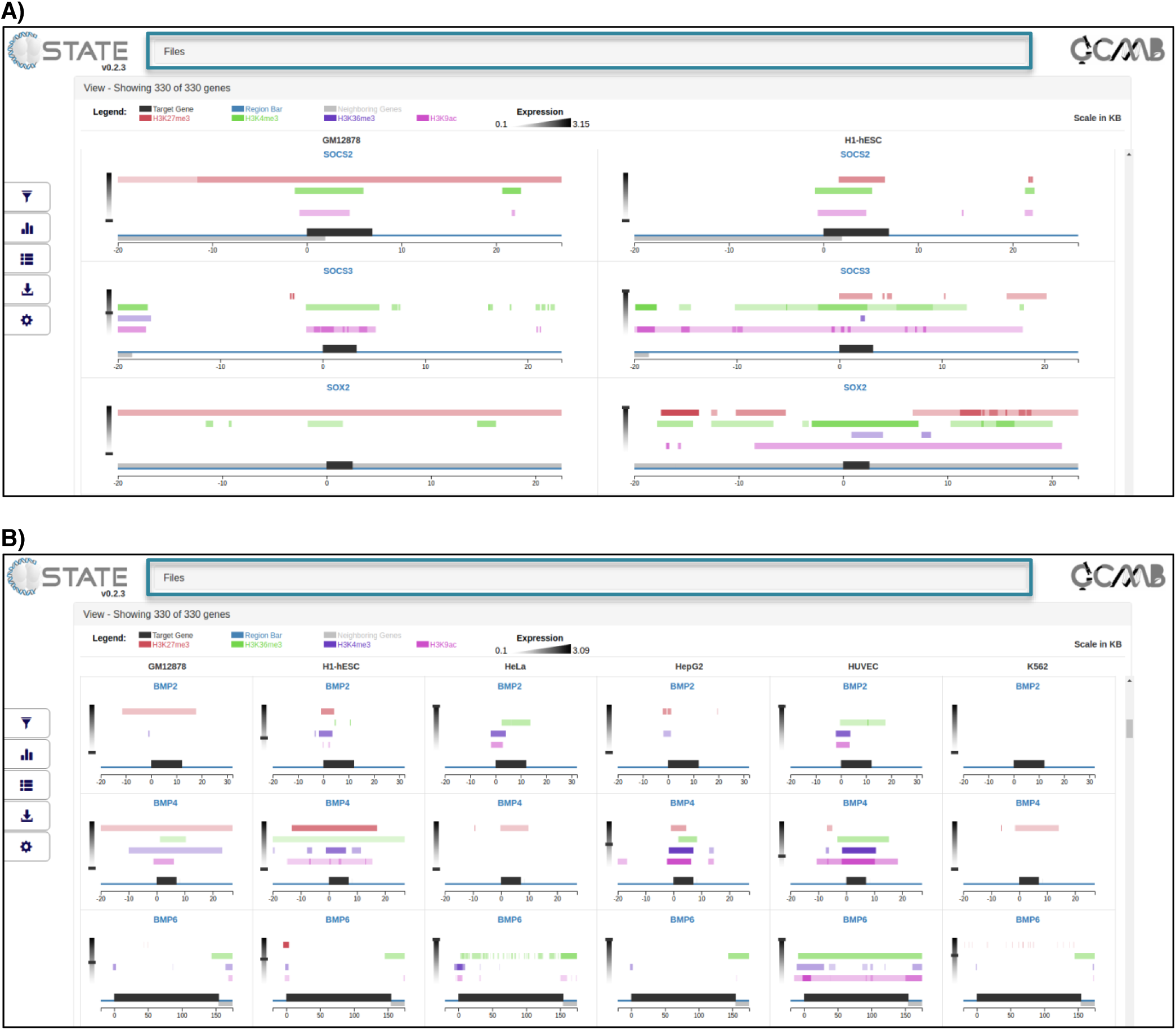
Open View accordion of C-State displaying gene data of **A)** two cell types and **B)** six cell types. Screenshot shows 3 of the 330 genes in the View pane. Blue rectangle indicates the folded Files accordion.

The region of interest is indicated by a scaled blue line with the target gene (indicated by its panel header) shown as a black bar on it while neighboring genes are depicted as grey bars in a strand specific manner. The scale is in kb (0 represents TSS) and specific to each gene in order to maintain visual similarity across all the genes, irrespective of size. Orientation of each gene is also taken into account for uniformity and enhanced visual comparison; all peaks from the data are calculated with respect to TSS, corrected for gene orientation, and plotted as shaded bars on multiple tracks of distinct colors above the gene. The opacity of the bars is a function of the peak intensity scores, which are displayed on mouse hover. Expression value of the gene in each cell type is displayed on a graded scale (default grayscale) on the side of the plot. The raw expression value is displayed on mouse hover.

#### Gene Modal view

The grid layout in the default display allows rapid browsing through all the genes in a list. However C-State also provides for gene-specific views across the chosen cell types and features. Clicking on any plot in the display opens a modal for the gene (Figure 4), where the data representing that gene in multiple cell-types is stacked vertically for closer inspection; the larger aspect ratio of a landscape layout allows focusing on an expanded viewpoint anywhere along the entire locus. The plots are interactive, and support panning as well as zooming (using either the mouse scroll or the zoom controls provided at the top of the modal), with the scale automatically adjusting to the zoom level. The zoom is linked to all the data tracks and cell types for a seamless comparison, and can be reset to the original state with the “Reset zoom” button. In addition, certain context specific information is displayed in the modal view such as the genomic coordinates and orientation of the gene (top left), exon information (alternating thick and thin bars to represent exons and introns respectively), peak intensity information and gene expression score or value (below cell type name).

**Figure 4:**
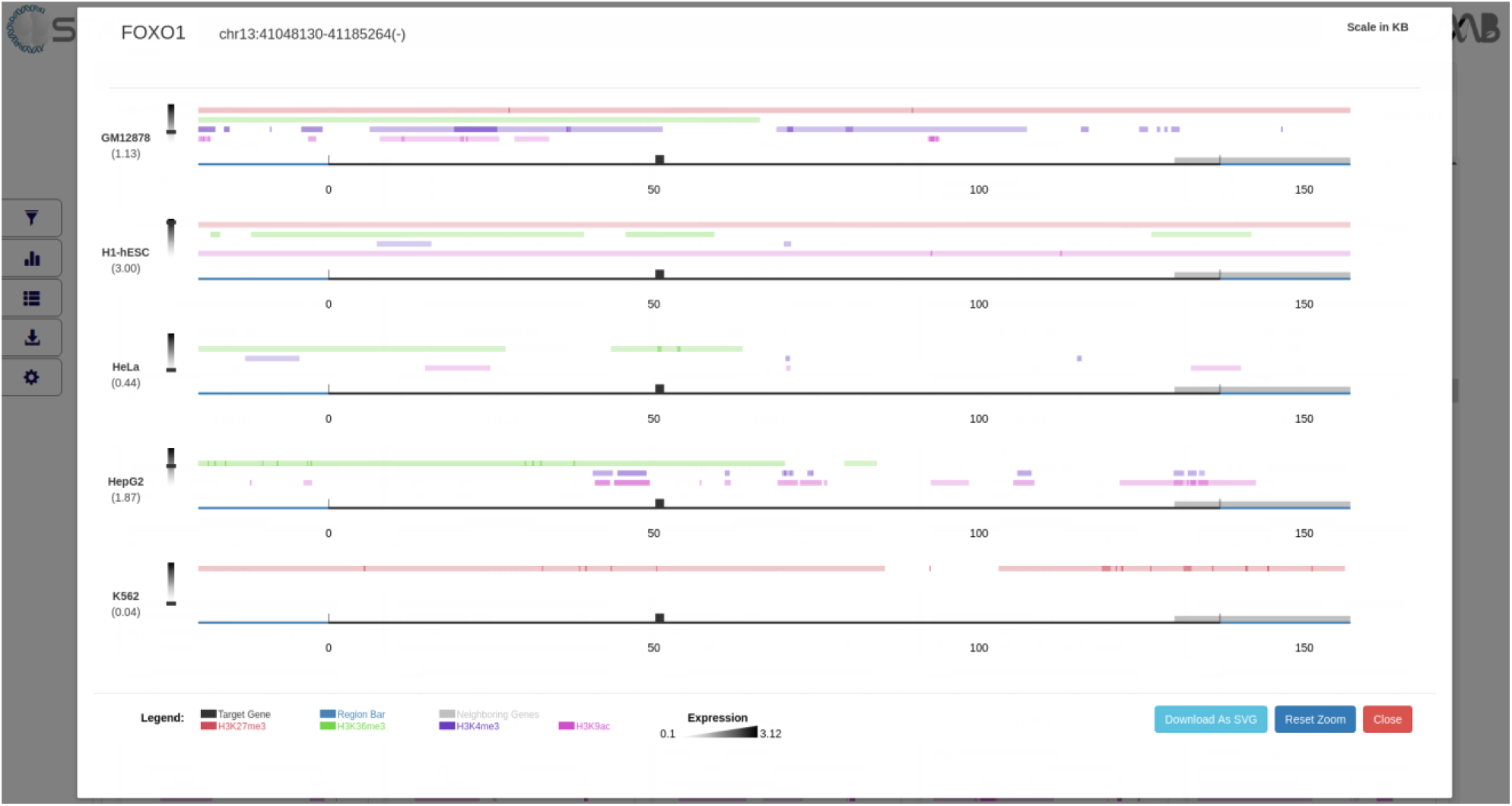
Expanded gene modal showing peak features (colored tracks) and expression values (heatmap scale on left, value in parenthesis) of a single gene (thick bars represent exons, thin bars introns) across 5 cell types stacked vertically.

## Case study

### Overview of datasets

Addressing biologically relevant questions often involves analyzing sets of genes belonging to particular pathways or regulating distinct cellular processes. However, extracting chromatin peak information of selected genes of interest from genome-wide datasets is a cumbersome task. The following use cases demonstrate the utility of C-State in the analysis of 16 epigenetic (4 histone marks across 4different cell types) and 4 RNA-Seq datasets from the ENCODE project [12]. We have focused on data from multiple human cell lines - K562, HeLa, and GM12878 - for comparison with H1 embryonic stem cells (H1- hESC) to examine changes in histone modification profiles. Whole genome ChIP-seq datasets are downloaded for H3K4me3 and H3K9ac (associated with gene activation), H3K36me3 (active transcription) and H3K27me3 (repression). The downloaded BED files are loaded directly into C-State as feature files in the Files accordion. The FPKM values of all genes derived from RNA-seq datasets of these cell types are loaded as expression data files (See Tutorial in the website for details and formats).

To identify enrichment patterns at a selected subset of genes in these differentiated versus pluripotent cell states, we created a list of ‘stemness’ genes potentially important for regulating the ES cell state from published datasets analyzing the hESC transcriptome [13] and pluripotency factor bound gene networks in hESCs [14]. A subset of 330 genes, shortlisted based on their change in expression profile upon ESC differentiation, is loaded into the Files accordion for comparative analysis. Gene expression patterns along with associated histone marks are analyzed across the group of 330 target genes and 20 KB of their flanking upstream and downstream regions.

The chained filtering application of C-State (Pattern Search module in Control Panel) allows instant identification of epigenetic patterns via simple queries as described below.

### Use case 1: Bivalent promoters in ESCs

Genes that have bivalent promoters (marked with both H3K27me3 and H3K4me3 within -5KB to +2KB of TSS) in ESCs can be identified using the “Feature Overlaps” Filter (Figure 5A) chained to a couple of “Feature Counts” Filters set for the absence of the other marks (Figure 5B). This returns just 13 (of 330) genes. Their individual gene modals can be examined from the View accordion or a list of details obtained from the Tables panel (“Show Filtered Genes only” box checked; Supplementary Figure S2). The bivalent gene names from the table can be directly copied to the clipboard for use in other applications.

**Figure 5:**
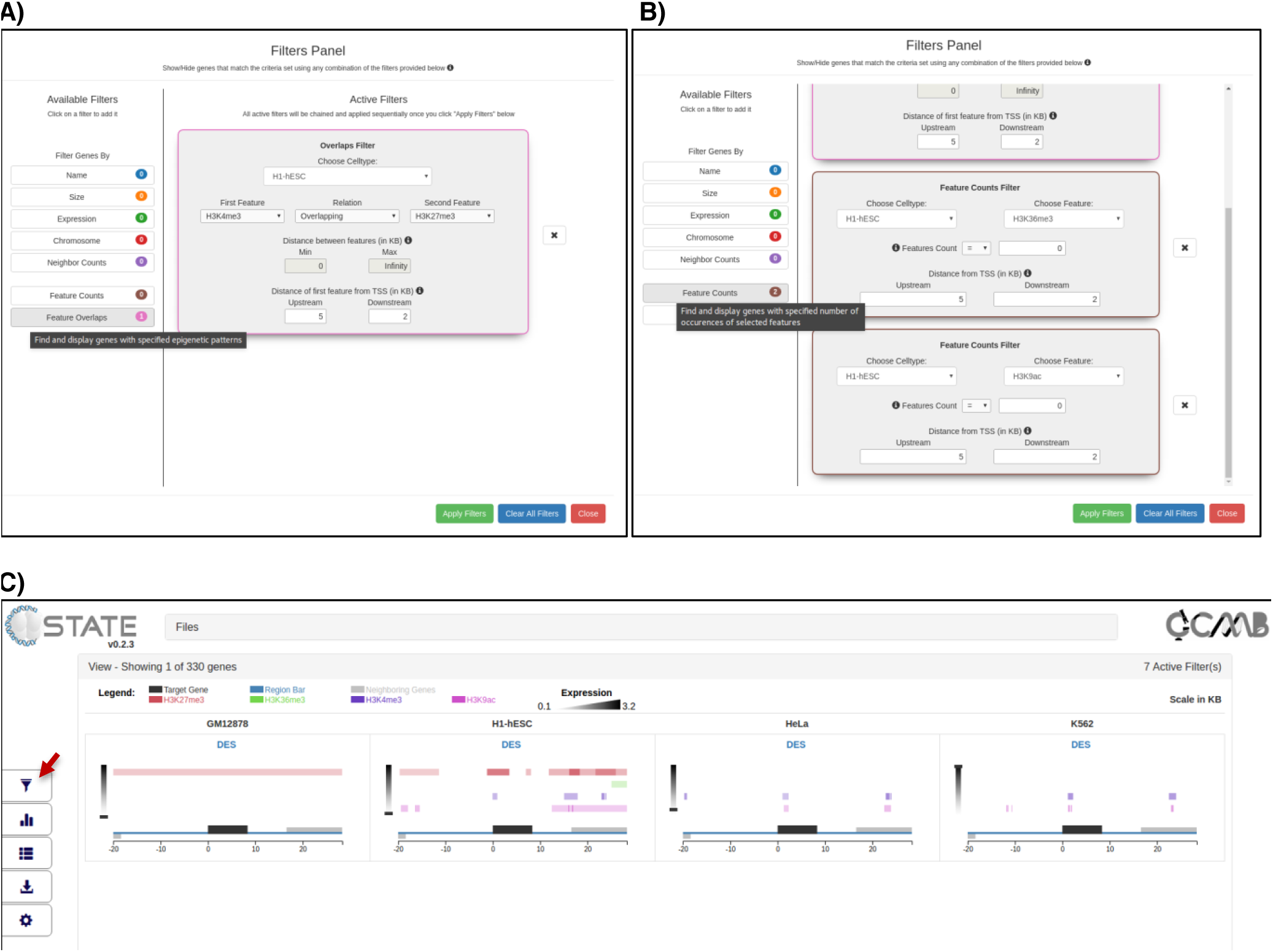
Pattern Search module (red arrow) in Control Panel showing **A)** Overlaps filter of C-State set to display genes with bivalently marked promoters (-5 kb to +2 kb of TSS) in ESCs and **B)** Two consecutive feature count filters added to the chain and set to further refine “clean” bivalent promoters (devoid of the other 2 marks, namely H3K36me3 and H3K9ac). **C)** View accordion of C-State with the filtered output - a single gene (DES) that is bivalently marked in H1-hESC cells at the TSS while carrying active marks in K562 (and HeLa) and enriched for repressive H3K27me3 in GM12878 cells.

To further identify genes where the ESC promoter bivalents resolve into a repressed chromatin state in one cell type (GM12878, Supplementary Figure S3A) and an actively marked one in another (K562, Supplementary Figure S3B), simply add the appropriate filters to the chain. Applying this chain of 7 filters instantly returns the muscle specific gene Desmin (DES), which has a bivalently marked promoter in ESCs that resolves into two distinct chromatin states in the other cell types (Figure 5C).

Plotting the average feature profile (Plots and Analysis key in Control Panel) in fact, reveals an increase in the average H3K27me3 enrichment around the TSS of genes in ESCs (Figure 6, 1^st^ row, 2^nd^ column) but not in other cell types. The distribution of other marks, however, remains the same across all cell types.

**Figure 6:**
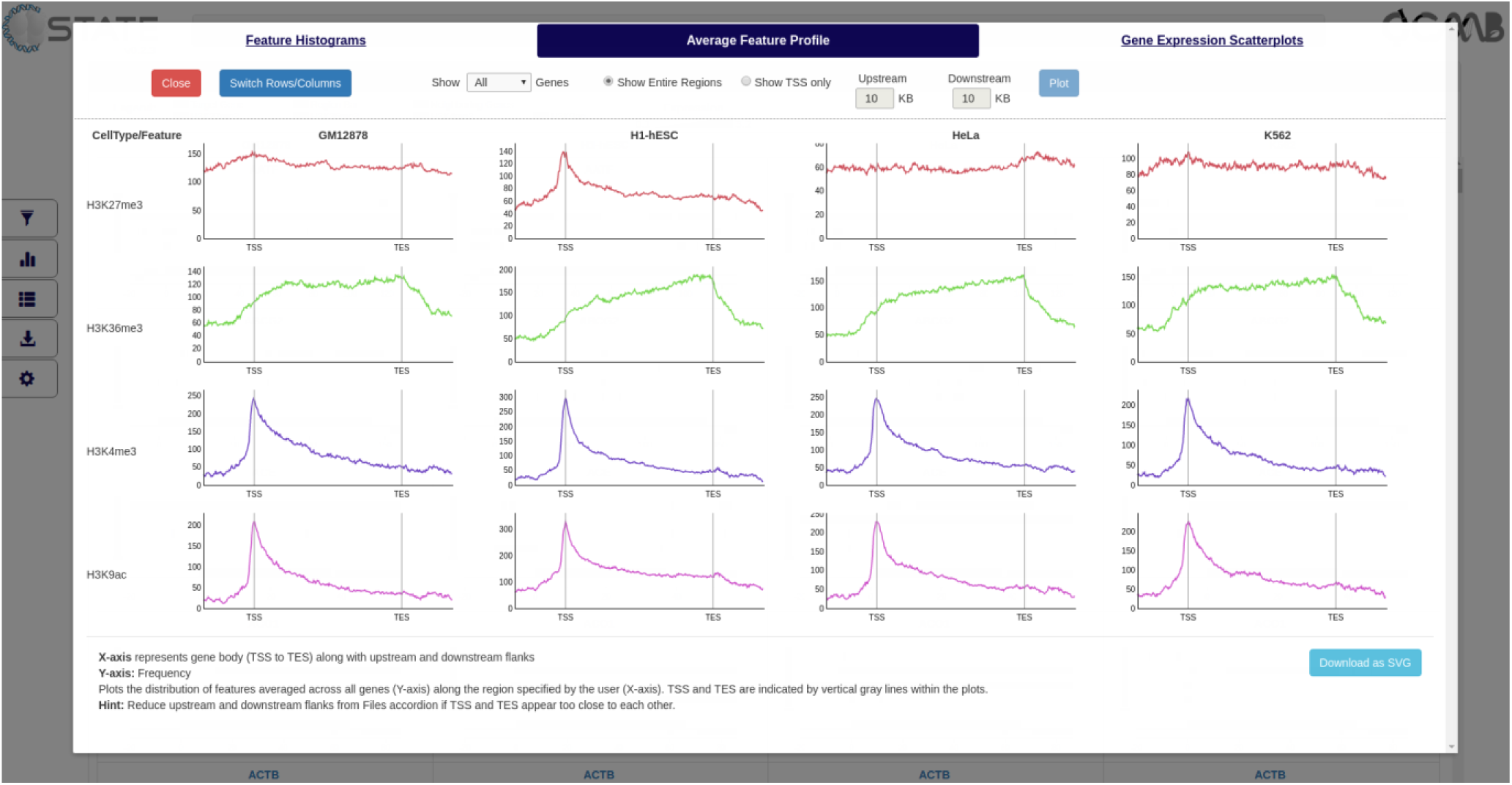
Plots and analysis module of C-State showing average feature profiles at gene bodies

### Use Case 2: Active Transcription in ESCs

To analyze change in histone mark distribution at actively transcribing genes in ESCs, select for those that are H3K36me3 enriched within 500 bp near exons in ESCs using the Pattern Search module - set the “Feature Overlaps” Filter for a maximum distance of 0.5 KB between H3K36me3 peak and an exon (Supplementary Figure S4, top). 97 genes are identified that match the above criteria in ESCs. Gene expression scatterplots (Plots and Analysis) show that these genes are indeed more expressed in H1-hESC cells as compared to the other three cell types (Supplementary Figure S5).

To further shortlist genes with high transcript levels in ESCs, add an “Expression” filter to the chain (Supplementary Figure S4, bottom). Set the cell type as H1-hESC, and set the minimum expression value to 3.2 (95th percentile of the loaded datasets, as depicted by the legend in the main view). This returns a list of 19 genes that are highly transcribed in ESCs. Descriptions of these 19 genes (Tables view with “Show Filtered Genes Only” ticked) indicate that they are developmentally important transcription factors. Chaining yet other filters to remove genes that are not expressed in other cell types returns target genes to focus on for epigenetic analysis, such as the pluripotency gene Nanog, the cardiac muscle alpha actin (Actc1) gene, and the brain-specific Notch signaling pathway gene, Notch3 (Supplementary Figure S6).

## Discussion

Change in chromatin state serves as a direct read-out for underlying regulatory mechanisms especially when correlated with change in gene expression status. Analyzing genes that behave in a similar manner (respond to cues with the same epigenetic profile) can help identify regulatory networks for cellular processes. In our efforts to examine the epigenetic status of a set of developmental genes across cell states, we realized that current tools do not offer the power of epigenetic analysis to the bench biologist. The number of genes needed to be analyzed is generally large (in hundreds) and putative regulatory elements may be present in proximal intergenic regions regulating gene expression by serving as binding sites to transcription factors and other chromatin-modulating proteins. Besides involving time consuming pre-processing and transfer of data to the available genome browsers servers, the examination of each of these genes individually is tedious and error-prone, and requires additional collating steps to comprehensively visualize and depict the observed patterns across genes.

The web based tools available for analyzing peaks of enrichment for epigenetic features or protein binding in genome-wide datasets provide limited scope for interaction and user-based querying for target regions. A few platform-specific packages achieve some balance for target selection and graphic visualization for pattern identification such as CHROMATRA [15], a plug-in specific for the Galaxy platform or PAVIS [16], which links to the UCSC browser and uses a chromosome-centric approach. Complete ChIP-seq analysis packages like CisGenome [17] and recently developed visualization tools such as VisPIG [4] and Epiviz [18] offer options for peak visualization but cannot directly screen and display user-defined patterns across multiple loci. Another challenge faced by biologists is that many of the current tools are command-line applications that rely on the computer- proficiency of the user or work only with an adequate understanding of the programming language they are written in. These include the bioconductor packages written in R [19, 20], Python packages such as seaborn and matplotlib that focus on graphical plotting [21], and MATLAB modules. Finally, most tools are specialized in a limited set of visualization tasks, making it an unstated requirement for users to be proficient in numerous related packages to solve all their plotting problems. The absence of a GUI with scope for flexible search and display options prevents biologists from understanding their data first hand and hence they rely on bioinformaticians to generate custom scripts and data-specific algorithms.

C-State is a handy tool for biologists as it does not require any programming skills to filter and analyze NGS data. C-State combines a module for the identification of enrichment patterns in context of gene transcription with an analysis module as well as a graphic visualization platform. It employs a simple, user-friendly GUI that enables easy identification of global as well as gene specific enrichment trends with no investments from the user other than providing a list of genes and the target datasets as the starting point. There are no imposed limits for target region or gene subset selection, nor on the number or complexity of the search patterns allowed. C-State supports simultaneous visualization of any number of genes and cell types or conditions with multiple tracks for each, limited only by the free memory available on the system (see FAQs on website). Designed as a web-app, C-State runs locally on the user’s system on all modern web browsers and is a standalone application that does not depend on any installations or data transfer via the internet. It also serves as a simple platform for data sharing and collaborative analysis without sharing raw data files. Additionally, C-State allows fast image generation to capture gene expression and epigenetic features changes at multiple loci in a single go. Table1highlights the features of C-State and compares their availability across other platforms.

C-State is thus ideal for both comparative search and analysis. Its key design choices enable rapid multi-gene visualization of a large number of features. Some tools offer an option of superimposing charts but this becomes cluttered when visualizing data from many conditions. To allow the simultaneous visualization of gene-specific changes across cell types, time courses or drug treatments, we opted for side-by-side views of a given gene. The “small multiples” based display module allows comparison across experimental conditions in a single shot grid view, a kind of superimposition that is not feasible in current genome browsers where each frame of view displays high resolution data only for a given locus. Visual space for genes utilizes the entire page while controls are folded away to the side (Control Panel) and the top (Files accordion). Superimposing the full screen Gene Modal over the View panel allows close inspection of a particular locus with multiple zoom options while scrolling down the View panel enables simultaneous gene to gene comparison. Users can use the search bar to quickly locate specific gene(s). Multiple names can be entered simultaneously; genes searched as a group are arranged together for easier comparison without scrolling through the entire set. Grouping of data tracks from each condition and converting features with respect to TSS based on gene orientation are other unique features that improve comparative visualization. The gene expression track within the view panel enables analysis of epigenetic changes in the context of their effect on gene expression, without having to open any other panel for checking cell-type specific transcription status.

C-State allows a large degree of customization. The user can restrict the features to be plotted by applying cut-offs in the settings pane on gene/peak attributes such as peak quality and size. The Feature Histograms from the Plots and Analysis panel can guide the user in choosing appropriate cut-offs. Similarly, the range of the gene expression scale can be adjusted based on the Gene Expression Scatterplots; the default range is calculated based on values within the 5^th^ and 95^th^ percentile in the loaded expression data. The genomic regions to be visualized around the target genes can be adjusted on the go using the flank selectors. The visual interface is fully customizable and the plot colors, track display, height and other features can be altered. Interactivity is enhanced with options such as mouse-over to display additional details such as peak size and score, gene name, size and neighbors, gene expression values and exon information. Every plot has adjustable controls such as organization of the sub-plot, axis range and definitions. Links within C-State facilitate navigation between the Views, Gene Modals and Tables by clicking on the gene name. Additional genomic details such as number of peaks and the gene expression status can be toggled from the table summary and the table can be sorted on any of the headers desired by the user.

While it is possible to upload several whole-genome datasets at once in other tools, data belonging to each condition has to be arranged manually for a meaningful comparison across conditions. Further, data of one condition or cell type is not considered or handled as a single group. This can often make data interpretation challenging and counter-intuitive if not handled computationally. Besides enabling epigenetic analysis in the context of cellular differentiation or treatment conditions, C-State can also be used for comparative epigenomics across disease states and cancer tissues or for any other genomic visualization such as single nucleotide polymorphisms (SNPs), mutation analysis in clinical and population genetics, genome annotations etc. C-State can help identify global trends in data. For example, the trend of distribution of a particular histone mark at gene bodies or peaking at a particular distance with respect to TSS can be identified from the Average Feature Profile plot. The marks and location can accordingly be set in the Filters panel to obtain selected genes that bear the same pattern. The Gene Expression Scatterplots can also similarly be informative regarding the cell types to choose to look for high (or low) gene expression.

C-State is designed for a wide audience of non-bioinformaticians but its open source and modular structure makes it easy for computer proficient users to extend its functionality. It embraces the philosophy of UNIX. provide a set of simple building blocks that can be combined to perform complex tasks. Currently, C-State exposes a simple API to toggle the display of genes, enabling the development of new filters with just a few lines of JavaScript. It supports many commonly used genome builds and we are in the process of adding support to other genome builds. There are a few limitations in the current version. Loading more than a thousand genes can slow down the app, depending on the available memory on the system. C-State supports limited input formats which can pose a restriction on users. Future versions will address these issues and incorporate more diverse features including additional pattern search modules and filters, and a flexible API for easier extensibility.

## Conclusions

C-State is a standalone application for epigenetic and gene expression analysis, providing an easy solution for experimental biologists to investigate epigenetic patterns without needing to know how to handle or parse big data. In addition to running locally on the user’s system, it can also be hosted centrally on an internal network to which multiple users can connect for visualizing and sharing their data. C-State’s searchable filtering and display modules are extensible and can handle data from any organism for a large number of genes and a flexible amount of intergenic regions. This is very useful for researchers to analyze novel or publicly available datasets in order to formulate new hypothesis for experimental testing, without investing in complicated programming or bioinformatics help. C-State also allows the incorporation of user-generated data with published datasets for rapid comparison and analysis and facilitates easy documentation of salient information via capture of high quality, publication ready images.

## List of abbreviations

API: Application programming interface
BED: Browser Extensible Data
ChIP: Chromatin immunoprecipitation
ESC: embryonic stem cell
FAQ: Frequently Asked Questions
GUI: Graphical User Interface
IGV: Integrative Genomics Viewer
MVVM: Model-View-View Model
NGS: Next Generation Sequencing
SVG: Scalable Vector Graphics
TSS: Transcription start site

## Declarations

### Availability of data and materials

C-State can be launched directly or downloaded for offline purposes from our website. Sample datasets, video tutorial, user manual and FAQs can be accessed at the C-State homepage. The source code of C-State is deposited in our github repository (https://github.com/RKMlab/c-state).

Project name: C-State Project home page:

http://www.ccmb.res.in/rakeshmishra/c-state/

Operating system(s): Platform independent

Programming language: HTML5/JavaScript

Other requirements: None

License: MIT

Any restrictions to use by non-academics: None.

### Author contributions

DTS and SS conceived and designed the study. SS conceptualized the application and performed the case study with comments and improvements contributed by JD and RKM. DTS developed and implemented the software. SS and DTS created the C-State website, tutorials and video documentation. SS drafted the manuscript with inputs from all authors. All authors read and approved the final manuscript.

## Acknowledgments

We thank Saurabh Gaur for the initial prototype of C-State. We are grateful to Hardik Gala and Gunjan Purohit for testing C-State and providing useful feedback. Saketh Saxena is acknowledged for the C-State website design.

## Ethics approval and consent to participate

Not applicable

## Consent for publication

Not applicable

## Additional Files

### Supplementary file 1: Supplementary figures

**S1:** Screenshot of the Filters Panel in the pattern search module showing all the 7 filters available (left) and the 4 filters opened in the Active Filters pane on the right. **S2:** Table view listing genes having bivalent domains at promoters in ESCs (13 of 330 genes). **S3:** Feature counts filters set to identify genes bivalent in ESCs that show a promoter (-5 Kb to +2 Kb of TSS) profile of A) H3K27me3 marks but no H3K4me3 enrichment in GM12878 cells and B) H3K4me3 peaks but no H3K27me3 enrichment in K562 cells. **S4:** Top: Feature Overlaps filter set to identify genes in ESCs that carry H3K36me3 enrichment at exons (within 0.5 Kb) indicating active transcription. Bottom: Gene Expression filter added to the chain to identify genes that additionally have high transcript levels. **S5:** Gene Expression scatterplot (Plots and Analysis) showing distribution of expression values of the 97 filtered genes obtained after setting the filter described in Figure 4, top. Many of these genes appear to be ESC-specific as they show medium to high expression in ESCs (boxed, column 2, X-axis represents expression in ESCs) compared to the other cell types. **S6:** View accordion displaying the genes filtered for high gene expression only in ESCs.

## Video demos are available from the C-State website

